# Characterization of genetically effective cells and EMS mutagenesis on the novel winter oil seed Pennycress (*Thlaspi arvense*)

**DOI:** 10.64898/2026.04.30.722012

**Authors:** Anthony Brusa, Ratan Chopra, Eva Serena Gjesvold, Matthew Ott, Liam Sullivan, Claire Chapman Biel, Shweta Jain, Carl Branch, Greta Rockstad, Krishan M. Rai, David Marks

**Affiliations:** University of Minnesota, Department of Agronomy and Plant Genetics, 1991 Upper Buford Circle, St Paul, MN, 55108; University of Minnesota, College of Biological Sciences, 1445 Gortner Ave, St Paul, MN 55108

**Keywords:** Pennycress, Ethyl methanesulfonate, Mutagenesis, Gene Index, Genetically Effective Cells, Intermediate Crop

## Abstract

Pennycress (*Thlaspi arvense* L.) is an intermediate winter oilseed crop that has only recently been domesticated for agronomic use. Improving agronomic traits requires sources of genetic variation, and mutagenesis is frequently used to help overcome the limitations of natural populations. We investigate the impact of Ethyl methanesulfonate (EMS) on genetically effective cells (GECs) to characterize the intra-individual genetic variation of EMS mutagenesis in pennycress. We identified that pennycress contains at least 4 GECs which, when treated with EMS, create unique mutations across different branches within the same individual plant. We then propagated the M2 plants for whole genome sequencing, providing extensive characterization of the EMS mutation profile and developing a gene index as a resource for future reverse genetic screenings.

**Article Summary:** Pennycress is an emerging winter oil seed crop in the American Midwest. Domestication efforts have advanced rapidly through a combination of genetic techniques. One of the most successful methods has been the use of a mutant gene index, a large collection of pennycress seed where new genetic variation has been created through Ethyl methanesulfonate (EMS). EMS mutations are not uniform however, and a single treated seed can have wide genetic variation within the resulting plant. We investigate the role of genetically effective cells on EMS variation, and present the full EMS population as a resource for further pennycress domestication efforts.

## Introduction

*Thlaspi arvense* L. (field pennycress or pennycress herein) collected from the wild has been targeted for domestication as a new intermediate crop that can be grown on tens of millions of acres in temperate regions between the harvest and sowing of traditional summer crops. Importantly, this new over wintering crop provides a number of ecosystem services. As a winter cover, pennycress can reduce soil erosion and it flowers early in the spring providing both honeybees and native pollinators with a new source of pollen and nectar. The ecosystem services of pennycress have been well characterized; including improvements in soil preservation (Cecchin et al., 2021), weed control (Johnson et al., 2015), pollinator services (Eberle et al., 2015; Evangelista et al., 2017), and water quality (Moore et al., 2020; Ott et al., 2019).Throughout its life cycle it can also take up and utilized excess nutrients applied the previous summer. This is especially desirable when following maize as even under the best circumstances maize only uses up to half the applied nitrogen, which can be up two hundred pounds per acre. This nitrogen typically either migrates to the ground waters, escapes in runoff polluting rivers, or is broken down into greenhouse gasses. By taking up excess nitrogen pennycress provides valuable ecosystem services to both the grower and broader community. However, the adoption of cover crops has been slow due the expense of sowing and terminating them in the field. Unlike cover crops, intermediate crops (such as pennycress) provide a harvestable product of value. Pennycress produces an oilseed with up 35% oil that is suitable for conversion to biofuels, bioplastics, renewable aviation fuel, and feedstock applications of pennycress meal (Hojilla-Evangelista et al., 2013; Markel et al., 2018; Moser et al., 2009).Thus, pennycress protects the land and provides the farmer with a new source of income during an otherwise unproductive season.

The initial wild pennycress isolates carried both desirable and weedy traits. As a winter annual pennycress can survive cold harsh winters, while yielding over 2000 lbs/acre of oilseeds in the mid to late spring. However, the yield can be greatly decreased by seedpod shatter during combine harvest. The high levels of the glucosinolate sinigrin decrease the value of the seed meal and the oil of wild pennycress contains high levels of toxic erucic acid. In addition, wild pennycress seeds have a thick antinutritional seed coat that allows pennycress to create a prolonged seed bank after harvest. Like Arabidopsis, pennycress is a diploid that is self-fertile. These features facilitate the use of classical chemical mutagenesis and gene editing to make homozygous recessive mutants with desirable traits. This has been demonstrated through both the screening of an EMS mutagenized population and through gene editing isolate new varieties of pennycress that lack undesirable traits (Chopra et al., 2020; Marks et al., 2021; Ott et al., 2021). This work was facilitated by the advent of next generation sequencing which was used to generate both a pennycress transcriptome and draft genome, which together defined nearly all of the gene space in pennycress. These analyses also showed that, similar to the closely related Arabidopsis, pennycress genes are mostly present as single copies (Chopra et al., 2018; McGinn et al., 2019). This led to the prediction that in pennycress it should be possible to isolate the same spectrum of mutants as found for Arabidopsis, and that it should be possible to identify causative candidate gene mutations in pennycress using Arabidopsis as a guide.

More recently, a revised annotated genome sequence was released in 2022 (Nunn et al., 2022) and is publicly available on NCBI (https://www.ncbi.nlm.nih.gov/assembly/GCA_911865555.2). This new genome sequence contains much more comprehensive annotations than previous assemblies, allowing for better identification and characterization of lines with important agronomic traits. This sequence was largely based on the natural accession MN106 that was isolated from a ditch near Coates, MN. Unlike previous versions of the genome, this new version makes use of a MN106 variant that had undergone 10 generations of single seed descent to reduce natural heterozygosity.

This newly sequenced MN106 variant was used to create an EMS mutagenized population. While the chemistry of EMS mutagenesis is quite well understood, the developmental biological process via which mutagenized cells gives rise to mutants in the next generation is not something that has been widely studied. In Arabidopsis, it is estimated that two to three genetically effective cells (GECs) in the dry seed embryo give rise to the gametes that form the next generation. During EMS mutagenesis of pregerminated seeds, each of the GEC accumulates distinct sets of mutations that are then passed onto the next generation. In Arabidopsis plant each GEC creates contiguous clones of cells from which seeds form on different branches or on different sides of the main branch of an individual (Irish & Sussex, 1992). This phenomenon was followed up on through phenotypic analyses, including Meinke and Sussex’s observations on the segregation of embryo lethal mutations in siliques on sectors of the main stem (Meinke & Sussex, 1979). In this case each of the embryo lethal mutants containing sectors included 1/3 to ½ of the circumference of the main stem, which is consistent with the presence of two to three GEC in the seed embryo. Similar investigations in *Lotus japonicus* estimated at least 3 genetically effective cells were present at mutagenesis (Perry et al., 2009). To investigate if this phenomenon holds true in pennycress, we used a novel approach by performing WGS on seedlings derived from the seeds collected from separate stems of individual M1 plants (plants derived from the original EMS treated seeds). This analysis provides new information on the potential number cells from mutagenized seeds that contribute to the M2 generation in pennycress. This information enables investigators to better predict the possible diversity generated by mutagenesis of seeds containing ungerminated.

To develop a new resource for pennycress domestication, EMS mutations in 857 M2 individuals were identified through WGS. This data was used to compile a gene index for use in reverse genetic screening. The improved annotation helped identify genic non-synonymous mutations. Extensive analyses were conducted on this dataset to determine mutation numbers and rates. As a proof of concept, the data supported a reverse genetic screen to find new mutants in the gene index population. The variant gene index data will enable researchers to use reverse genetics to find lines with specific mutations in a gene of interest, potentially speeding up the discovery of new agronomic traits. To streamline this process, we developed a workflow that allows researchers to identify and filter mutants for a specific gene and obtain seed from a known, genotyped mutant pennycress plant in less than six months

## Materials and Methods

### 1 EMS mutagenesis

Genetic variation for the mutant gene index was generated using EMS (ethyl methanesulfonate) mutagenesis on MN106ref seeds. Fifty thousand seeds were placed on a shaker and imbibed with distilled water overnight. These seeds were then treated with 0.2% EMS in a 0.1M sodium phosphate buffer (pH7.0) for 18 hours. EMS was deactivated by immersion in 0.1M sodium thiosulfate (pH7.3) for 20 minutes. Mutagenized seeds were washed in distilled water and then soaked in a 50 µM GA_4+7_ solution overnight.

Mutagenized seeds were planted in a field plot measuring 5ft x 50 ft using the broadcasting method, and M1 plants were vernalized and flowered in the field. Seeds from more than 1,000 randomly selected M1 plants were harvested. Seed from the M2 individuals was planted in the Fall of 2020 and 2021, harvested the following spring, and provided the M3 seed, which serves as the MGI’s long-term genetic resource.

### 2 DNA isolation from M1 and M2 shoots and DNA sequencing

To enable the creation of a mutant gene index and to track segregation of alleles post-EMS treatments in branches of M1 plants and M2 bulk seed lots, we collected DNA from the seedlings and single plants, respectively. To understand GECs in pennycress, we collected seeds from 7-10 unique stems/branches of M1 plants. Approximately 20 seeds from three different branches and one main stem of an M1 plant were planted on filter paper, and DNA was collected from 15-day-old seedlings using the methods described below.

To build the mutant library, M2 seeds from individual M1 plants harvested in the fields was be planted on filter paper with water in a sandwich bag. Seeds were allowed to germinate for 7-10 days and then transferred to a cold chamber for vernalization. After 21 days of cold treatment, seedlings were transferred to soil in 2-inch containers for establishment and phenotyping. We grew wild-type (MN106-Ref) seeds for comparison using the same method. We collected leaf tissue from the individual which had the most robust pod fill in order to produce viable seeds for future experiments.

Approximately five young leaves from a plant of interest, or ∼20 seedlings from each line of interest were collected using forceps in a 1.5 microfuge tube. These tubes were immersed in liquid nitrogen before being transferred to -80 °C for later extraction. DNA was isolated using either the theGenCatch TM Plant Genomic DNA Purification Kit (http://www.epochlifescience.com/) or the Qiagen DNEasy Plant Mini Kit (httpe://www.qiagen.com) with some modifications to produce high-quality DNA. Briefly, frozen tissue was crushed using a microfuge tube-compatible pestle, and PX1 buffer containing RNase was be added and homogenized by vortexing. Tubes were incubated at 65 °C for 10 minutes, and then gently mixed after 5 minutes. The PX2 buffer was be added to the tubes and incubated on ice for 15 minutes. After incubation, tubes were centrifuged at 15000 rpm, and the supernatant was removed for further processing on the DNA column. The final elution (55 µl) was performed using DNase/RNase-free water. We quantified DNA samples using nanodrop (https://www.thermofisher.com/). The highest-quality samples were sent to the Joint Genome Institute for genome resequencing on an Illumina Novaseq at 15x coverage under Proposal ID: 506821.

### 3 Bioinformatics

Sequence data of pennycress accessions were processed using a standard bioinformatics workflow. Raw reads were processed using Trimmomatic (Bolger et al., 2014) to remove low quality reads before being aligned through bwa-mem (Li & Durbin, 2009). Alignment was performed against the pennycress v2.0 genome (Nunn et al, 2022). Post-alignment processing was performed using Samtools and Picard tools to remove PCR duplicates, assign read groups, and for sorting. Variant calling was be performed on each of the mutant aligned files separately with the GATK tool Haplotype Genotyper (McKenna et al., 2010) for both SNPs and indels. We predicted the effects of mutations in the coding regions and regulatory sequences using SNPEff (Cingolani et al., 2012).This required construction of an SNPEff database built on the v2.0 pennycress genome, and the VCF file from GATK was used as input to generate an annotated VCF file. Annotations included amino acid changes based on open reading frame, addition or removal of stop codons, variant impact, and location of the variant relative to known genes. The VCF was filtered to remove intergenic variants and the remaining variants were exported to an Excel file for ease of use. One additional filtration step was performed to create a tab containing only non-synonymous variants.

### 4 Candidate searches

We developed a pipeline for processing and evaluating lines and used it to identify two different examples of successful reverse genetic screenings using the MGI. Candidate genes for MGI validation were identified through ongoing projects and were part of well documented pathways, supported by literature search (Liljegren et al., 2004; Sagasser et al., 2002). All variants within these genes were identified through the MGI and considered for evaluation. Amino acid sequence alignments were performed using Clustal Omega (Sievers et al., 2011) and functional annotations for the genes were obtained through Uniprot (https://www.uniprot.org/). Lines with early stop codons or mutations in functional regions were targeted for evaluation. Seed from candidate lines were germinated in a 50 µM GA_4+7_ solution. Seedlings were then vernalized for 3 weeks, transplanted into flats, and transplanted into 72-well flats in a growth chamber. At the 4 leaf stage leaf tissue was collected and genotyped using a competitive allele specific PCR (PACE, https://3crbio.com/) to identify homozygous mutants. These individuals were then grown to maturity, allowed to self, and evaluated for the mutant phenotype against MN106 as a control. Transparent testa mutants were phenotyped visually due to the stark difference between mutant (yellow) and wildtype (black) seed coat. Shatter resistance (IND) was phenotyped using a 5N force gauge to measure the force required to break open a mature seed pod. 30 measurements were taken per individual plant, and statistical significance was determined using comparison of means (T-test).

## Results

### 1 Impact of genetically effective cells (GEC) on EMS mutagenesis

An analysis of mutations across the main stem and three branches of individual M1 plants showed a complex combination of both shared and private mutations (figure 1).In the individual Sec_7 we found that 31,638 out of 42,700 mutations (73.9%) were only present in seeds collected from a single branch. The remaining mutations were present in the seeds collected from at least two branches, with only 2,981 mutations (6.9%) present in all 4 branches that we sequenced. Similar results were observed in the other two individuals; SEC_8 had 25,844 out of 39,791 (64.9%) mutations present only in seeds collected from single branches and 3,240 (8.14%) found in seeds collected from all branches, while SEC_10 had 19,835 mutations out of 31,751 (62.4%) present in seeds collected from single branches and 1,512 (4.76%) found in seeds collected from all branches. Across all three individuals analyzed we found that private mutations, those found exclusively in a single branch, accounted for a majority of mutations present.

**Figure 1.**
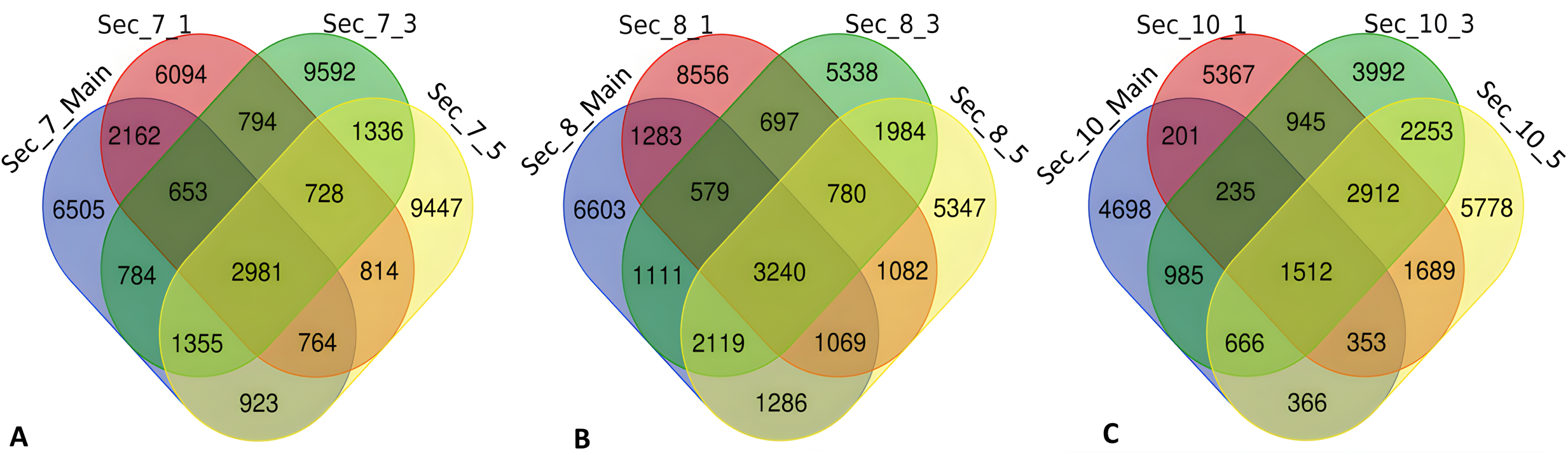
A comparison central stem and three branches show that EMS mutagenesis creates a mix shared and private mutations within branches of the same individual. Private mutations made up the majority of mutations across Sec_7 (A), Sec_8 (B), and Sec_10 (C).

Similar results were also found when sequencing M2 individuals which were derived from seed taken from differing branches of the same M1 individual (figure 2). Two siblings descended from the M1 plant 201-M2-55 had 9,221 shared mutations, with the sibling plants containing 9,668 and 8,301 private mutations resulting in a roughly even split of private and shared mutations across siblings (32.7% shared, with siblings having 34.3% and 33% private). 2020-M2-142 showed a similar pattern, with 6,610 (34.2%) of mutations shared between siblings and 12,800 (46.85%) and 7,911 (29%) mutations being private. Seedlings derived from 2020-M2-933 showed a different pattern with only 282 (3.83%) mutations being shared, while the vast majority of mutations (3,938 or 53.6% and 3,127 or 42.6%) were found only in one of the siblings.

**Figure 2.**
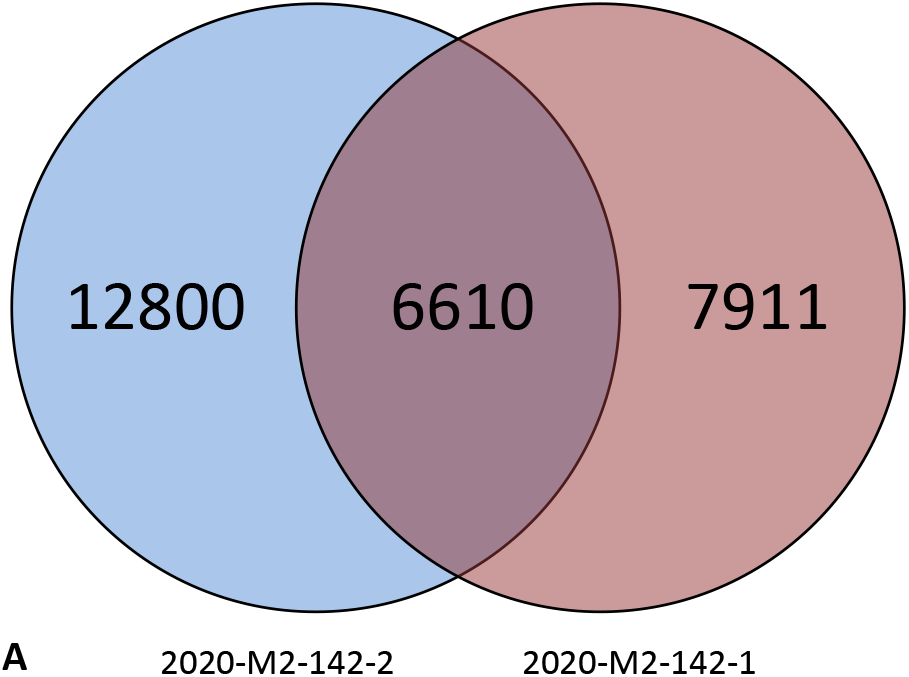
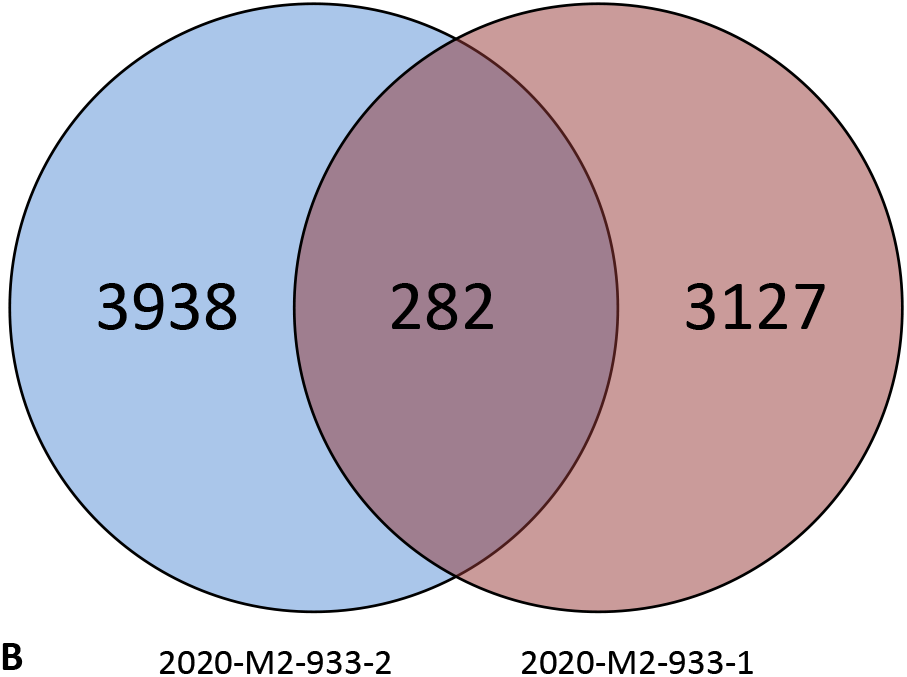
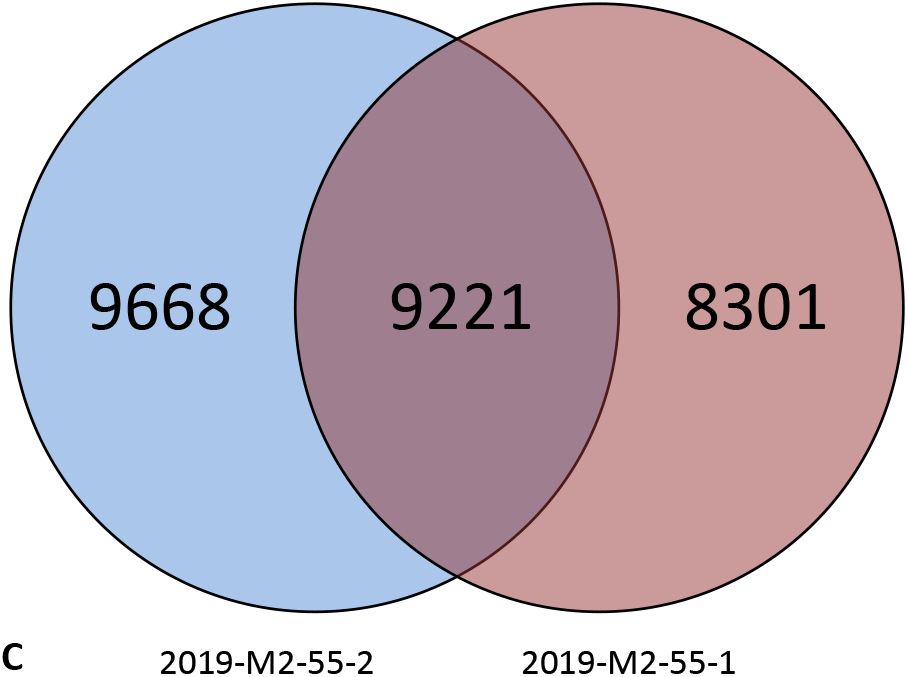
A comparison of shared and private SNPs in M2 individuals from three lines; 2020-M2-142 (A), 2020-M2-933 (B) and 2019-M2-55 (C)

These results are consistent with a limited number of GECs, but at least four, in the pennycress embryo that mostly but not exclusively produce contiguous clones in the aerial shoot to create the large numbers of stem specific private mutations. However, the data is also consistent with the hypothesis that after mutagenesis and during subsequent cell divisions that there is some mixing of the original progenitors of the distinct populations of mutagenized EMC cells. This latter phenomenon would account for the shared mutations observed between the various stems whose seeds served as the source for the WGS analyses. This phenomenon has not previously been reported and is a departure from the results observed in *Arabidopsis*.

### 2 Characterization of M2 EMS mutations

To generate a genome-wide resource for pennycress mutants, 857 independent M2 EMS-mutagenized plants, each derived from a single M1 plant were subjected to WGS. This represents only a small fraction of the diversity created by EMS in these populations, but it ensured that each M2 individual contained a distinct non-overlapping set of mutations. For the final index the nature and distribution of both genic and non-genic EMS-induced mutations were characterized (Table 1). 3,871,244 total mutations were observed in genic (and non-genic???) regions with 3,676,458 (95.0%) of mutations being single nucleotide variations (SNVs). Deletions and insertions represented 117333 (3.0%) and 77453 (2.0%) of mutations respectively. EMS mutagenesis preferentially causes transitions, which was reflected in a transition rate of 80.42% among SNVs. Indels represented only 5% of total variation, with 1bp (2.33%) and 2 bp (1.11%) indels accounting for the majority of these variants. Large indels (6+ base pairs) represented 0.83% of all variants.

**Table 1:** Mutation overview of EMS mutagenized lines.

Total mutations
| Features | Number |  |
| --- | --- | --- |
| Single Nucleotide Polymorphisms (SNPs) | 3676458 |  |
| Deletions | 117333 |  |
| Insertions | 77453 |  |
| Total Number of Mutations | 3871244 |  |
| SNP mutations | Number | Percent of Total Variation |
| A > T | 54025 | 1.40 |
| T > A | 57480 | 1.48 |
| C > T | 1562272 | 40.36 |
| G > A | 1551287 | 40.07 |
| A > G | 98684 | 2.55 |
| T > C | 98405 | 2.54 |
| G > C | 24975 | 0.65 |
| C > G | 25919 | 0.67 |
| A > C | 35281 | 0.91 |
| C > A | 66123 | 1.71 |
| T > G | 36525 | 0.94 |
| G > T | 65482 | 1.69 |

Further evaluation (Table 2) showed that, among mutations in genic regions, 3,635,065 (93.9%) were found outside of exons (in non-genic??? Non-coding regions) and 73,213 (1.9%) variants were synonymous changes within exons. A total of 162,963 (4.2%) variants were located within exons and found to be non-synonymous. The vast majority of these, 150345 (92.2%) were missense mutations which cause single amino acid changes. Higher impact mutations were also observed; 8,160 (5%) early stop codons, 6,591 (4%) changes in splice sites, 4,084 (2.5%) frame shift mutations, and 304 (0.1%) loss of start codons. Out of 29,236 annotated genes, 88.27% had some form of unique mutation, 21.96% had a premature stop codon, and 8.69% had a frame shift mutation. Additional details, including an index of non-synonymous mutations, are available in supplemental table S1.

**Table 2:**
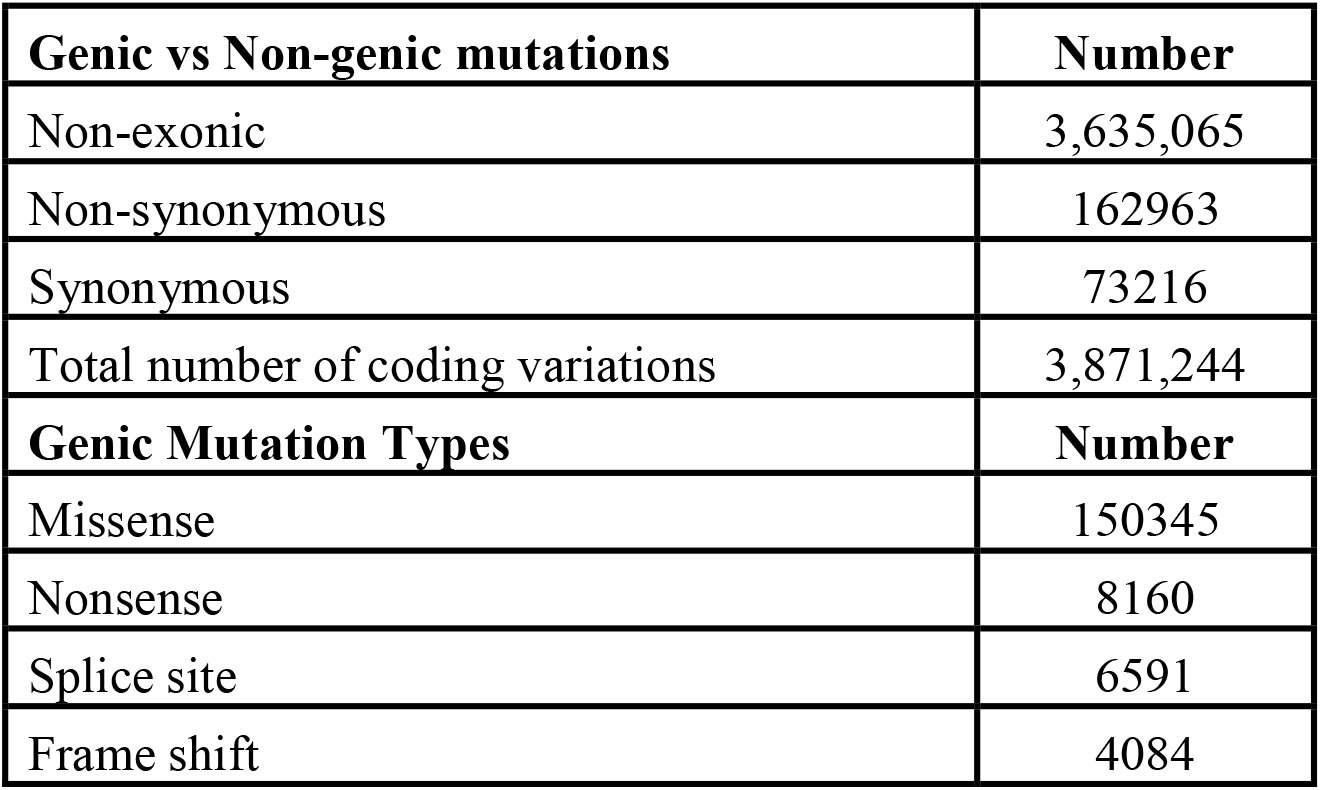
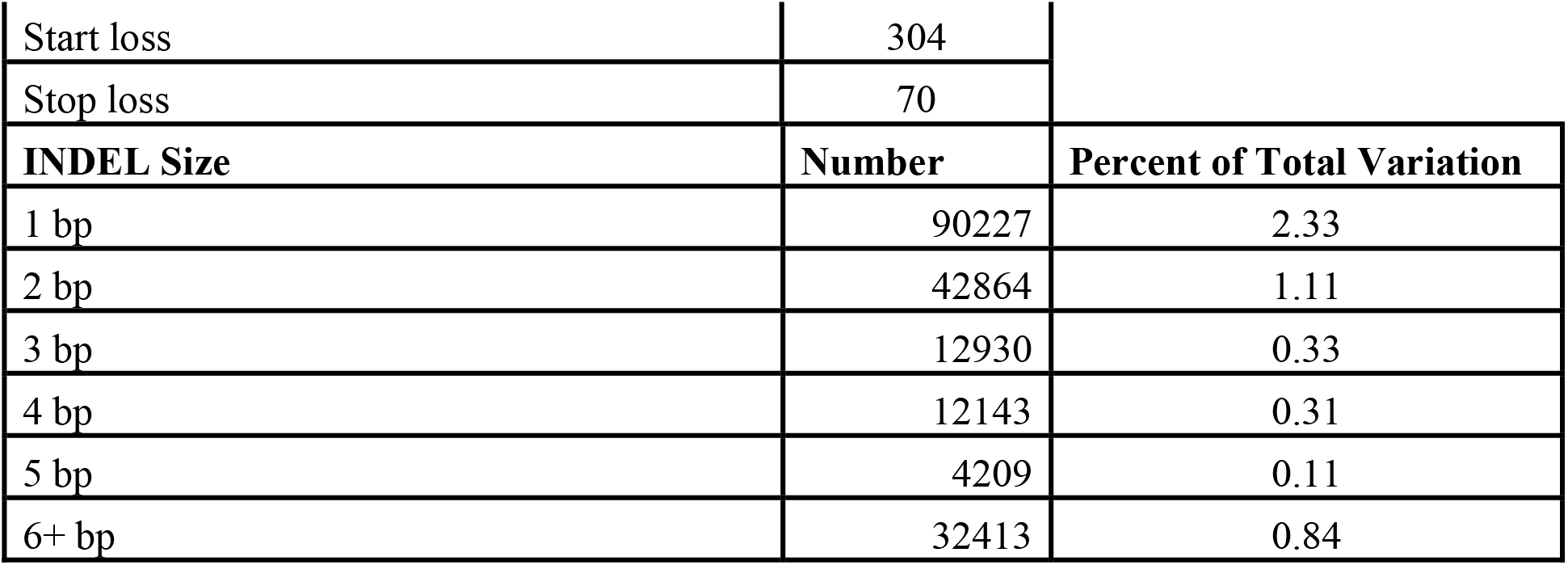
Feature overview of EMS mutagenized lines.

| Genic vs Non-genic mutations | Number |
| --- | --- |
| Non-exonic | 3,635,065 |
| Non-synonymous | 162963 |
| Synonymous | 73216 |
| Total number of coding variations | 3,871,244 |
| Genic Mutation Types | Number |
| Missense | 150345 |
| Nonsense | 8160 |
| Splice site | 6591 |
| Frame shift | 4084 |

|  |  |  |
| --- | --- | --- |
| Start loss | 304 |  |
| Stop loss | 70 |  |
| **INDEL Size** | **Number** | **Percent of Total Variation** |
| 1 bp | 90227 | 2.33 |
| 2 bp | 42864 | 1.11 |
| 3 bp | 12930 | 0.33 |
| 4 bp | 12143 | 0.31 |
| 5 bp | 4209 | 0.11 |
| 6+ bp | 32413 | 0.84 |

We also selected random mutant sites from the individuals in the library to make molecular markers to verify the presence of predicted mutations in the M3 and M4 generations. For the M3 generation, we found that 36 of 41 mutants sites tested were present in the later generations (supplemental S2). For the M4 generation, we found that 24 of 30 sites tested could be confirmed as segregating (supplemental S3). These results demonstrate that the majority of the alleles identified at M2 generation are still present in M3 and M4 generations. However, the success rate will be lower in more advanced generations due to some mutant alleles getting fixed (1/4 of the induced mutations per generation as the original M1 GEC would have been heterozygous for any particular induced mutation) or lost due to segregation (1/4 of the induced mutations per generation). Overall, after many generations if there was no selection pressure, at most ½ of the original mutations would be expected to still be in the population, while the other ½ would segregate away before being fixed. Finally, it is likely that many fixed homozygous or heterozygous mutations also could be lost due to fitness issues. These fitness issues may be caused by the alleles themselves, linked alleles, or fixed alleles elsewhere in the genome. Importantly, growing the plants in a field environment allowed only plants to be selected that are sufficiently fit to survive the challenging winter conditions of the Upper Midwestern United States. Many of the induced mutations associated with unfit plants would be lost.

## 3 Validation of EMS populations for use in reverse genetics

### 3.1 Reverse genetic screen for pennycress seed coat mutants

The identification of transparent testa mutants with lighter seed coat color is a straightforward and easy trait to phenotype and to follow. Therefore, it was selected as our initial target for which to identify mutants in the MGI via reverse genetic screening. Previous work had shown that that pennycress seed coat color was altered by mutations in genes involved in the production of condensed seed coat tannins as was also seen with *Arabidopsis* (Chopra et al., 2018). Thus, several genes in the pathway, including the transcription factors TT2 and TT8 along with the genes TT6, TT7, BAN, TT12, which encode pathway enzymes, were used as targets for a reverse genetic screen. For this screen sequenced lines were identified that carried a mutation in one of these genes (Table 3). Seed packets containing pooled seeds from these MGI lines were screened for the presence of lighter colors seeds, which is indicative of a defect in pathway or regulatory genes (Figure 3). Typically, strong loss of function mutations in either TT2 or TT8 result in the formation of yellow seeds, while mutations in pathway genes result in various shades of reddish to tan colored seeds. Interestingly, one of the three tt2 mutants displayed tan seeds instead of yellow. However, the mutation in this mutant was a nonsense mutation that truncated only the last three amino acid residues in the peptide. Thus, this likely was the result of the creation of a partial loss of function (lof) allele. All of the other mutants had phenotypes similar to those previously seen for strong loss of function mutations in these genes.

**Table 3:** Characteristics of MGI lines used in reverse screening for seed coat mutants.

| Gene | Line | CHROM | POS | REF | ALT | Aa substitution | Phenotype |
| --- | --- | --- | --- | --- | --- | --- | --- |
| TT2 | 2019-M2-207 | 7 | 46787912 | C | T | Arg260* | tan |
| TT2 | 2020-M2-050 | 7 | 46787753 | G | A | Asp207Asn | yellow |
| TT2 | 2020-M2-136 | 7 | 46787409 | C | T | Ser92Phe | yellow |
| TT8 | 2019-M2-071 | 6 | 50472264 | AAG | - | Glu230del | yellow |
| TT8 | 2019-M2-138 | 6 | 50470953 | C | T | Ser122Phe | yellow |
| TT8 | 2019-M2-078 | 6 | 50472678 | G | A | Trp317* | yellow |
| TT8 | 2020-M2-832 | 6 | 50471441 | G | A | Gly133Arg | yellow |
| TT6 | 2020-M2-076 | 5 | 4728376 | G | A | Gln325* | tan |
| TT6 | 2020-M2-404 | 5 | 4728244 | C | T | Ala369Thr | tan |
| TT7 | 2020-M2-032 | 6 | 67293391 | G | A | Thr247Ile | tan |
| TT7 | 2020-M2-624 | 6 | 67292562 | C | T | Trp469* | tan |
| BAN | 2020-M2-117 | 5 | 67793133 | C | T | Ala182Thr | tan |
| TT12 | 2020-M2-029 | 5 | 1610688 | G | A | His215Tyr | tan |
| TT12 | 2020-M2-865 | 5 | 1609716 | C | T | Gly422Glu | tan |

**Table 4:** Characteristics of MGI lines used in reverse screening for shatter resistance.

| Gene | Line | CHROM | POS | REF | ALT | Aa substitution |
| --- | --- | --- | --- | --- | --- | --- |
| IND | 2020-M2-918 | 6 | 69868 | C | T | Tarvense_00017:p.Gly53Asp |
| IND | 2020_M2_245 | 6 | 70001 | G | A | Tarvense_00017:p.Leu9Phe |

**Figure 3.**
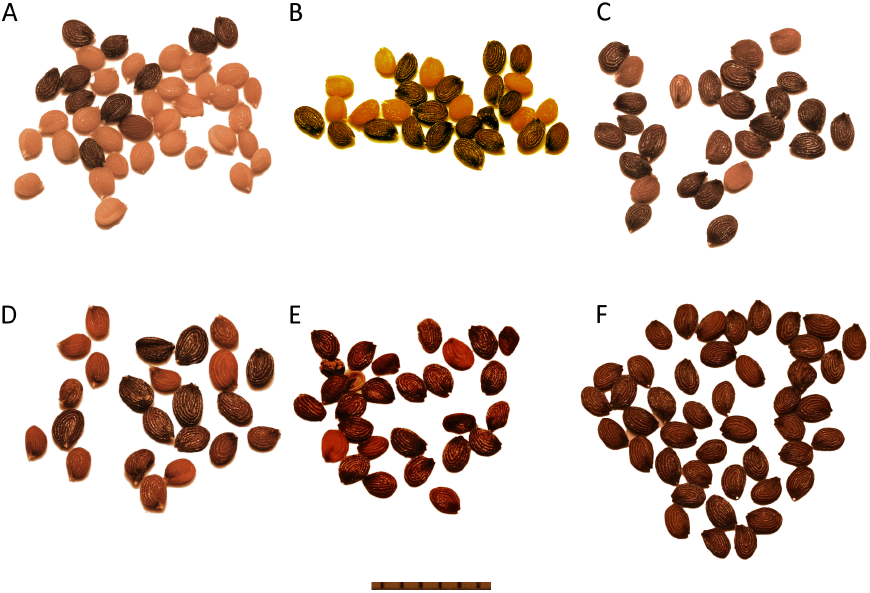
Images of seeds from selected pooled MGI line seeds. MGI lines were screened for the presence of non-synonymous mutations in genes encoding peptides required for seed coat condensed tannin formation. Shown are images from seed packets containing a mix of wild-type like and lighter colored mutant seeds. A) 2020-M2-207 tt2; B) 2019-M2-138 tt8; C) 2020-M2-404 tt6; D) 2020-M2-624 tt7; E) 2020-M2-029-tt12; F) MN106 parental control.

Overall, 34 lines were identified carrying mutations in either regulatory or pathway genes. Within this population 14 lines or 41% of the lines exhibited phenotypes consistent with carrying a mutation in gene required for normal seed coat development. Not unexpectedly, the best predictor of a phenotype was the presence of a nonsense mutation. Four of five lines with gene stops displayed altered seed coat coloration. We cannot rule out the possibility that the fifth mutation might also have caused an altered seed coat, but this particular mutation was likely lost due to segregation in the generation that was used to create the mutant pool. However, this loss should be a rare event as the seed pools typically were created by combining M3 seeds from at least eight M2 individuals to capture as many of the original EMS induced mutations as possible. In any case, this screen validates the utility of the gene index population as a new source of mutants that can be identified via reverse genetic screens.

### 3.2 Reverse genetic screen for seedpod shatter resistance

IND (indehiscent) is transcription factor involved in the differentiation of cells responsible for fruit dehiscence (Liljegren et al., 2004). It is an important gene for domesticated pennycress; a partial loss of function in IND results in phenotype with reduced seed pod shatter, expressed as a higher force required to spate the halves of a seed pod (Chopra et al., 2020). Four mutations in the MGI were identified within the IND gene, two of which demonstrated a reduced shatter phenotype (Table 3). These mutations correspond to an amino acid substitution of Gly53Asp and Leu9Phe.

Plants were grown in a growth chamber alongside the reference line MN106. Reduced shatter trait was assessed by using a 5N force gauge to measure the force required to open seed pods, and 30 pods were measured per individual plant. Both amino acid substitutions were accompanied by significant increase in the required force for opening seed pods (Figure 4, P<0.0001).

**Figure 4.**
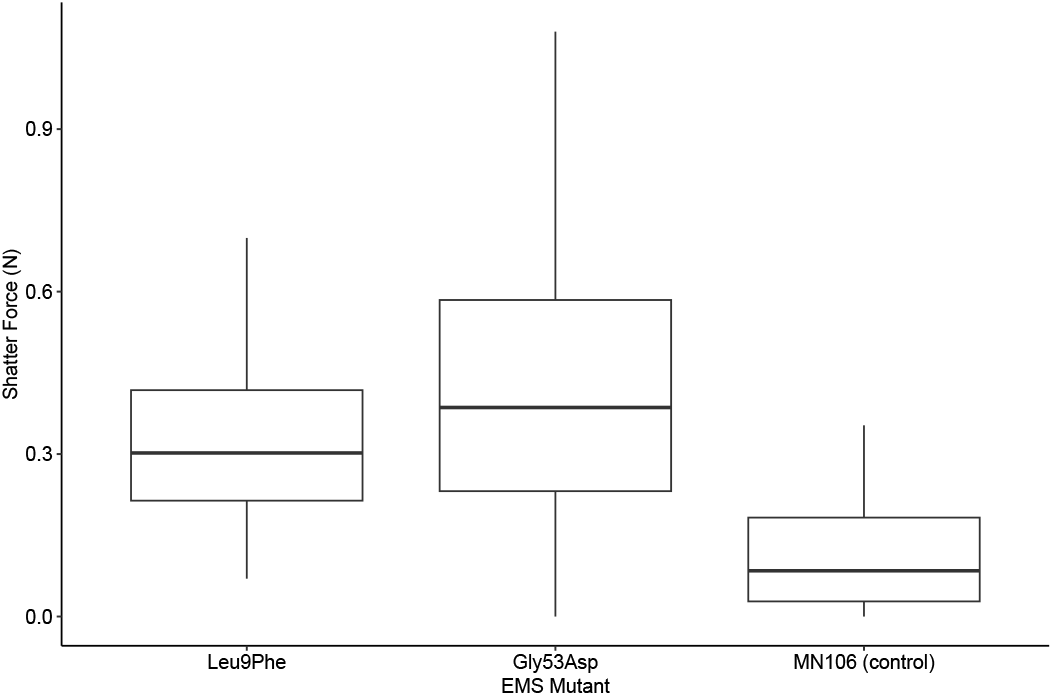
Two single amino acid substitutions (Leu9Phe and Gly53Asp) in the gene IND show a significant increase in force required to shatter seed pod compared to wildtype MN106.

## 4 Discussion

### GEC and EMS mutagenesis

According to classical plant genetics many dicot embryonic apical meristems are composed of a two cell layer tunica in which cells propagate via anticlinal divisions to maintain the two layers as distinct entities. The larger corpus domain lies beneath the tunica. Cells in this core region undergo both periclinal and anticlinal divisions to generate the bulk of the cells. An interesting feature of most dicots including members of the *Brassicaceae* is that a small number of cells in the second layer of the embryonic tunica give rise to both female and male gametic tissue. This is estimated to be two to three cells. Importantly, each of these co-called genetically effective cells (GEC) form clones of cells that are maintained as distinct vertical sectors up one side of a plant or the other. Within each sector, the clone of cells contains the progenitors for both male and female gametes. This greatly simplifies the ability to conduct chemical mutagenesis on seeds from self-compatible plants as a mutation that is created in one of the GEC is transmitted to the next generation via a classic one homozygous mutant to two heterozygous to one homozygous wild-type segregation ratio. Thus, it is possible to identify homozygous mutations in seeds that form on one of the distinct sectors of the plant.

To test this classic interpretation of plant development, M2 seeds were collected from the main stem and the three main side shoots of three M1 different plants derived from EMS mutagenized seeds. EMS randomly creates heterozygous mutations in the diploid sets of chromosomes in the GEC. The expectation was that whole genome sequencing of seeds from the side shoots would reveal the presence of distinct pools of mutation. With the caveat that adjacent side stems would share some mutations. Further, it was predicted that seeds from that main stem would contain a unique set along with additional sets of mutations present in each of the distinct side stems. As shown in Figure 1 this prediction was largely satisfied. Within each dataset, four distinct pools of mutations were identified. This supports the notion that pennycress contains at least four distinct GEC in the embryonic embryo of the seed. The sharing of mutations between stems on the same plant shows that each stem was derived from more than one distinct sector of cells, and thereby contains mutations derived from multiple GEC. Same for the main stem as having been derived from one main GEC but also sharing mutations with seeds derived from the side shoots. This indicates that that the progeny of the embryonic GEC create both shared and non-shared cell populations that give rise to the next generation of male and female gametes. The discrete pools of mutations indicate the present of sectors. But the sharing of mutations between putative M1 sectors indicates the presence of some intermingling of tissues derived from individual GEC cells. To further confirm this, we also sequenced two independent M2 derived from a single M1 plant, and we identified several unique sites in each of the M2 lines (figure 2). Two of these comparisons (2019-M2-55-2 and 2020-M2-142-2) showed roughly equal amounts of private and shared mutations between the siblings, although 2020-M2-933-2 showed a much stronger divergence with very few shared mutations between the two individuals observed. In all three cases, the initial effect of GEC conveying different of mutations across branches impacted the availability of such mutations in the following generation.

### Implications of GEC for reverse genetic screening

The results of the GEC work demonstrate that pooling across multiple M2 individuals is important for the creation of the most complete mutant resource. By pooling across multiple sibling plants the full spectrum of EMS induced mutations can be captured, providing the maximum potential for reverse genetic screenings. The tradeoff for this is that many individuals will not have the mutation of interest, requiring a broader screening pool to obtain mutant seedlings with the desired mutation. In our validation work above we found that a 72-well flat reliably provided enough homozygous mutants for reverse genetic screening. Through the use of optimal growth chamber settings and competitive fluorescence genotyping we were able to obtain mutant seed in approximately 4 months. Allowing for 2 generations of seed increase, this means that the turnaround from initial investigation to having enough mutant seed for agronomic evaluation of a trait can be completed in a single year.

### Reverse Genetic screen

Previously, we reported on using a pennycress EMS population for reverse genetic mutant screen (Chopra et al., 2018). While we were able to recover a few mutants, the population in our 2018 paper presented some challenges due to the parental pennycress plants used for that work having only recently been isolated from the wild. WGS sequencing analyses showed that this parental population was mixed with over 131,000 natural SNPs (Dorn et al., 2015). These extra SNPs complicated the use of reverse genetics as many or most of the alleles in the population were natural and not induced by EMS. To address this, the initial wild population (designated MN106) was subjected to single seed descent over 10 generations. The progeny from the tenth generation were used to create the EMS population described in this report. The finding that over 80% of the SNPs identified in this current populations were expected G to A or C to T transitions (Table 1) supports the notion that vast majority of the SNPs in this population were induced by the EMS treatment. Thus, this population will be superior in performing reverse genetic screens compared to the previous resource.

As a proof of concept, we were able to scan the current MGI mutant sequences to identify several classes of mutants that carry important domestication traits. These included new alleles of transparent testa mutants. These mutants have value as they typically have reduced indigestible seed coat fiber making the residual meal available after oil extraction more valuable as an animal feed supplement. In addition, these mutants typically display reduced dormancy to allow for better field establishment in the fall, and importantly, the mutations can reduce the creation of long lived pennycress seed banks that are created by wild-type dark seeds. The second class carried mutations in the IND gene is that is required for the formation of the indehiscent zone in the seed pod (Liljegren et al., 2004). In wild-type plants this zone promotes the breakage or opening of the seed pods to release the seeds. In mutants with a reduced indehiscent zone more force is needed to break the pods. This greatly improves yield by reduces loss due to wind induced pod breakage while still allowing pods to break open in the combines used for harvest.

While many of the MGI mutants could be recreated by new gene editing technologies, this WGS sequenced MGI population still has value as it enables discovery of rare alleles that can be crucial for agronomic traits and plant fitness. For one, the population allows ready access in one generation to many potential mutants that would require more time to recreate using gene editing. Second, it is likely that many desirable alleles will be partial loss of function alleles that are readily created by EMS mutagenesis but far more challenging for gene editing. For example, complete loss of function mutations in IND result in seedpods that are highly resistant to shatter – even in combine harvesting where unbroken seedpods are disposed of out the back of the machine. The IND mutants identified in this report still show some seedpod breakage, indicating that the causative mutations are partial rather than complete loss of function mutations.

Future domestication of pennycress will involve traits obtained from a variety of sources. Traditional breeding continues to make gains in complex polygenic traits, while forward genetic screenings of both natural and EMS induced variation are being deployed to identify the basis of new traits. Reverse genetics is an important complement to these approaches, with CRISPR allowing for precise knockout of candidate genes and optimized reverse genetics workflows enabling rapid isolation, evaluation, and seed increase of new mutant lines for downstream agronomic study.

## 5 Data Availability

Sequence data from this project can be found on Joint Genome Institute under Proposal ID: 506821 as part of award DOI: 10.46936/10.25585/60001359

## 6 Acknowledgements

The authors would like to thank Brett Heim and Dr. Katherine Frels for their help with field propagation of the lines used in this work. We would also like to thank the Don Wyse for his support of pennycress domestication efforts over the last 10 years at University of Minnesota.

## Figures

Attached seperately

## Author Contributions

Anthony Brusa: Methodology, Investigation, Visualization, Validation, Writing – Original Draft, Writing – Review and Editing

Ratan Chopra: Conceptualization, Methodology, Investigation, Funding Acquisition, Data Curation, Writing – Review and Editing

Eva Serena Gjesvold: Investigation, Data Curation

Matthew Ott: Investigation, Data Curation

Liam Sullivan: Investigation, Data Curation

Claire Chapman Biel: Investigation, Data Curation

Shweta Jain: Investigation, Data Curation

Carl Branch: Investigation, Data Curation

Greta Rockstad: Investigation, Data Curation

Krishan M. Rai: Investigation, Data Curation

David Marks: Conceptualization, Investigation, Validation, Supervision, Funding Acquisition, Writing – Review and Editing

## Legends for Supporting Information

Supplemental Table 1: Additional mutation characteristics of 857 EMS mutagenized pennycress lines, including an index of non-synonymous mutations

Supplemental Table 2: 36 out 41 randomly chosen loci showed segregation for the library alleles in the M3 generation

Supplemental Table 3: 24 out 30 randomly chosen loci showed segregation for the library alleles in the M4 generation

